# Cold-induced switch in direction of chloroplast relocation occurs independently of changes in endogenous phototropin expression

**DOI:** 10.1101/544361

**Authors:** Yuta Fujii, Yuka Ogasawara, Yamato Takahashi, Momoko Sakata, Saori Tamura, Yutaka Kodama

## Abstract

When exposed to fluctuating light intensity, chloroplasts move towards weak light (accumulation response), and away from strong light (avoidance response). In addition, cold treatment (5°C) induces the avoidance response even under weak light conditions (cold-avoidance response). These three responses are mediated by the phototropin (phot), which is a blue-light photoreceptor and has also been found to act as a thermosensory protein that perceives temperature variation. Our previous report indicated that cold-induced changes in phot biochemical activity initiate the cold-avoidance response. In this study, we further explored the induction mechanism of the cold-avoidance response in the liverwort *Marchantia polymorpha* and examined the relationship between changes in the amount of phot and the induction of the cold-avoidance response. The switch between the accumulation and avoidance responses occurs at a so-called ‘transitional’ light intensity. Our physiological experiments revealed that a cold-mediated decrease in the transitional light intensity leads to the induction of the cold-avoidance response. While artificial overexpression of phot decreased the transitional light intensity as much as cold treatment did, the amount of endogenous phot remained unchanged by cold treatment in wild-type *M. polymorpha*. Taken together, these findings show that the cold-avoidance response is initiated by a cold-mediated reduction of the transitional light intensity, independent of the amount of endogenous phot. This study provides a clue to understand the mechanism underlying the switch in direction of chloroplast relocation in response to light and temperature.

## Introduction

Plants sense ambient light and temperature to adapt to an ever-changing environment. In response to the light and temperature changes, the intracellular chloroplast position is precisely controlled. Under weak light conditions (e.g., under canopy), chloroplasts move towards light to increase light capture for photosynthetic efficiency (accumulation response) [1]. Under strong light conditions (e.g., under direct sunlight), chloroplasts move away from light to avoid photodamage (avoidance response) [2]. Chloroplast position is also affected by temperature [3], notably cold. Under warm conditions, weak light induces the accumulation response; however, under cold conditions (5°C) the same light level induces chloroplast avoidance (Fig 1). This movement referred to as the cold-avoidance (or cold-positioning) response, has been studied in several plant species, including the moss *Funaria hygrometrica* [3], evergreen ferns (e.g., *Adiantum capillus-veneris*) [4], the liverwort *Marchantia polymorpha* [5] and the flowering plant *Arabidopsis thaliana* [6]. In *M. polymorpha*, the cold-avoidance response contributes to the reduction of photoinhibition under cold conditions [6].

**Fig 1.**
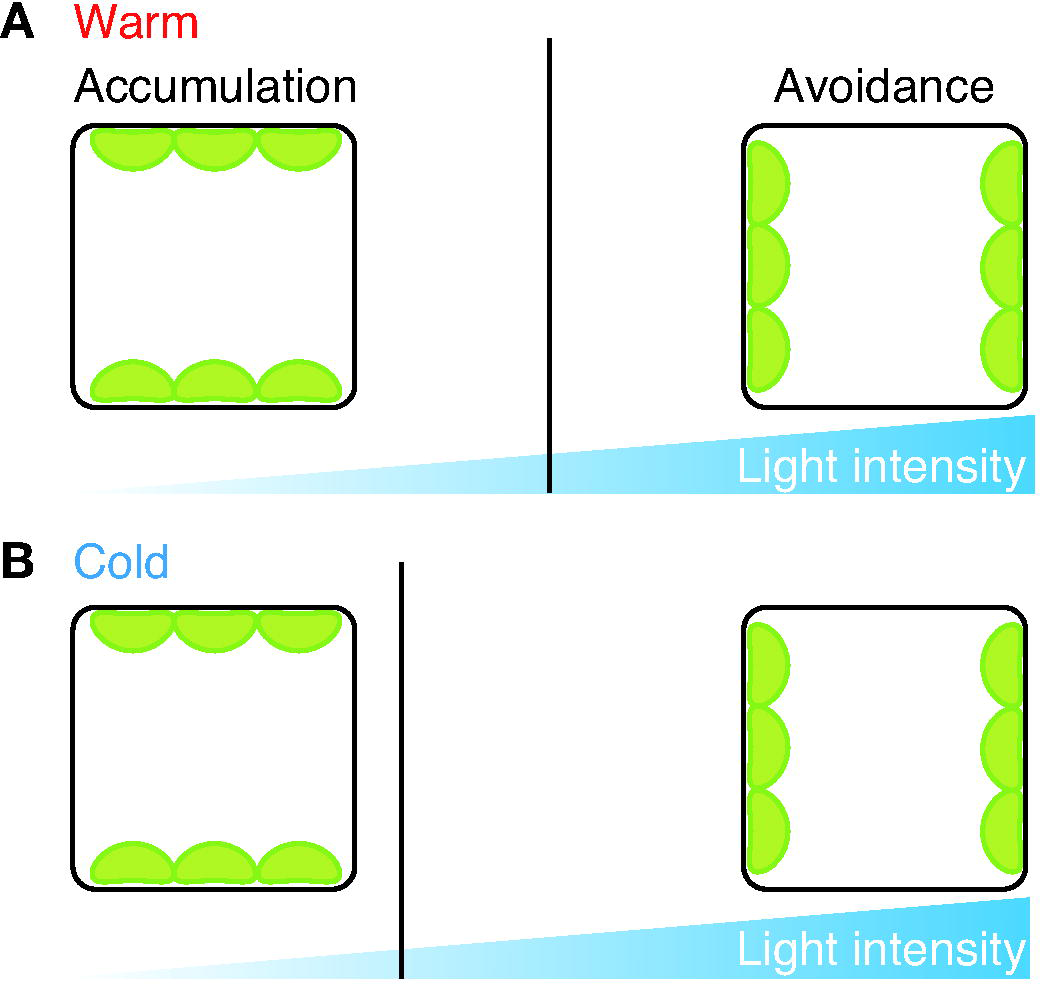
Schematic diagram showing the transitional light intensity of the accumulation and avoidance responses. Solid vertical lines indicate points of the transitional light intensity under warm (A; 22°C) and cold (B; 5°C) conditions.

Earlier studies showed that the accumulation, avoidance and cold-avoidance responses are all mediated by the blue-light (BL) photoreceptor phototropin (phot) [4,6–9]. Phot has two photosensory light, oxygen, or voltage (LOV) domains (LOV1 and LOV2) at its N-terminal region and a serine/threonine kinase domain at its C-terminal region. Phot perceives BL via its LOV domains and mediates the change of chloroplast position upon photoactivation of the LOV domains and kinase-mediated autophosphorylation [10]. A decrease in ambient temperature prolongs the lifetime of the photoactivated LOV2 domain, resulting in an increase in autophosphorylation [6]. The cold-induced enhancement of the photochemical and biochemical activities of phot (i.e., a prolonged lifetime of photoactivated LOV2 and increased autophosphorylation status) is essential for induction of the cold-avoidance response [6], and further study is required to fully understand this induction.

The switch in chloroplast behavior between the accumulation and avoidance responses occurs at their ‘transitional’ light intensity (Fig 1A). In the cold-avoidance response, because the chloroplast moves away from even weak BL (wBL) under cold conditions [6], it is hypothesized that the cold-avoidance response is induced by a cold-mediated reduction of the transitional light intensity (Fig 1B). However, this hypothesis remains to be tested. Artificial overexpression of phot has been suggested to decrease the transitional light intensity in *A. capillus-veneris* and *A. thaliana* [11, 12]. In addition, expression levels of endogenous phot are affected by both light [7,8,13–16] and temperature [16] conditions. Based on these findings, we postulate that a variation in the endogenous amount of phot may be involved in the induction of the cold-avoidance response.

The goals of this study were two-fold: first, to determine if a cold-mediated reduction of the transitional light intensity initiates the cold-avoidance response, and second, to determine if changes in phot amounts play a role in the induction of the cold-avoidance response. *M. polymorpha* was chosen for this study because it has a single copy of the gene for phot (Mpphot) that can mediate the accumulation, avoidance, and cold-avoidance responses [6, 17].

## Materials and Methods

### Plasmid construction

To construct a plasmid to generate the complementation line of Mpphot-Citrine (35S::Mpphot-Citrine/Mp*phot^KO^*), we used the Gateway cloning system (Invitrogen) and the binary vector pMpGWB306 [18], which includes *Citrine* (a gene for yellow fluorescent protein) for fusion at the 3 -end of the target gene under the control of the L cauliflower mosaic virus (CaMV) 35S promoter. The LR reaction (Invitrogen) was performed with a mixture of pMpGWB306 and the donor plasmid pDONR207-MpPHOT [19], and the resulting plasmid pMpGWB306-MpPHOT was used for transformation of *M. polymorpha*.

### Plant materials and growth conditions

Thalli of *M. polymorpha* were asexually cultured on half-strength B5 medium under 75 µmol photons m^-2^ s^-1^ continuous white light (FL40SW, NEC Corporation). Light intensity was measured by using a LI-250A light meter (LI-COR Biosciences). The Mp*PHOT* knockout line (Mp*phot^KO^*) was generated and kindly provided by Komatsu et al. [17]. In this study, transformation of *M. polymorpha* was performed by the G-AgarTrap method [20, 21], which is an *Agrobacterium*-mediated transformation method used for gemmalings. To generate the transformants, we used the binary vector pMpGWB106-MpPHOT [19] for Mpphot-Citrine overexpression lines (35S::Mpphot-Citrine/WT), and pMpGWB306-MpPHOT for the complementation line of Mpphot (35S::Mpphot-Citrine/Mp*phot^KO^*). The Citrine-expressing line (35S::Citrine/WT) was generated in our previous study [20]. G2 gemmalings were used throughout.

### Observation of chloroplast positioning

One-day-old gemmalings were incubated in temperature-controlled incubators (IJ100, Yamato Scientific Co., LTD) and light conditions were adjusted using blue light-emitting diodes (LEDs; OptoSupply Limited) and a LI-250A light meter. To produce polarized weak BL, 25 µmol photons m^-2^ s^-1^ of the blue LED was used as weak BL [5, 6], and it was filtered with a polarizing filter (HN38, Polaroid Corp.). Chlorophyll fluorescence images were captured using a fluorescence microscope MZ16F (Leica Microsystems) equipped with a DP73 digital camera (Olympus) and quantification of chloroplast positioning was performed using the P/A ratio method with ImageJ software, as previously described [5]. Briefly, fluorescence intensity was measured at 30 points (0.625 µm each, equivalent to 1 pixel) along the anticlinal cell walls (A) and 30 points (39.1 µm^2^ each, equivalent to 10 × 10 pixels) along the periclinal cell walls (P). This procedure was repeated five times, and then the P/A ratio was calculated as a mean of the five P/A values. The standard deviation was calculated using Excel software (Microsoft).

### Immunoblot analysis

Immunoblot analysis was performed as previously described [6]. Briefly, four-day-old gemmalings were collected in liquid nitrogen and kept at -80°C until use. The frozen plants were homogenized in SDS buffer (2% SDS, 10% glycerol, 5% mercaptoethanol, and 0.25 M Tris-HCl pH6.8) or Nonidet buffer (0.5% Nonidet P-40, 0.5M EDTA, 0.15 M NaCl, and 0.01 M Tris-HCl pH7.5) using a mortar and pestle and then boiled at 95°C for 5 min. The samples were centrifuged at 14,000 g for 10 min, and the supernatants used. Total protein amount was quantified by Bradford-XL assay (APRO Science). Fifteen µg of protein was subjected to SDS-PAGE with 8% polyacrylamide gels and transferred to a polyvinylidene difluoride (PVDF) membrane [6]. MpPHOT proteins were detected with rabbit antibody raised against the custom-made synthetic peptide (C+VDERAPPSKGSAKE) (Eurofins Genomics) and horseradish peroxidase-conjugated anti-rabbit IgG (Thermo Scientific). Both antibodies were used at dilution of 1:2000. For histone detection, anti-Histone H3 antibody (Abcam) at dilution of 1:5000 and horseradish peroxide-conjugated anti-rabbit IgG (Thermo Scientific) at dilution of 1:2000 were used. Chemiluminescent signal was detected using ECL-Select (GE Healthcare) and Light Capture (ATTO). For the image of Rubisco large subunit (RBCL), fifteen µg protein was subjected to SDS-PAGE with 8% polyacrylamide gels and stained with conventional Coomassie brilliant blue (CBB) method.

## Results

### Chloroplasts avoid weak blue light (wBL) under cold conditions

Using temperature-regulated microscopy with a microbeam irradiation system, we previously reported that under cold conditions (5°C), chloroplasts moved away from wBL that induced the accumulation response at 22°C [6]. However, in the study, physiological evidence of wBL avoidance under cold conditions was limited to microbeam analysis [6]. Here, we further analyzed whether chloroplasts move away from wBL under cold conditions.

In *M. polymorpha*, both cold and BL were required to elicit the cold-avoidance response [6]. Building on the result, to explore whether the cold-avoidance response is due to avoidance from wBL under cold conditions, we tested the effects of changing light direction. First, to induce the cold-avoidance response, wild-type (WT) cells were incubated under 25 µmol photons m^-2^ s^-1^ of wBL (BL25) with vertical irradiation at 5°C (Fig 2A). In WT cells, chloroplasts were positioned along the anticlinal cell walls as the cold-avoidance response (Fig 2B). When the light direction was changed from vertical to horizontal (Fig 2A), the chloroplasts along the anticlinal cell walls moved to the periclinal cell walls (Fig 2B), indicating chloroplast avoidance from BL25. To further confirm the results, we employed polarized wBL, because the vertically vibrating polarized wBL could induce chloroplast localization along the anticlinal wall by the accumulation response in *A. capillus-veneris* [22, 23]. When 10-day-old sporeling cells of *M. polymorpha* were irradiated with vertically vibrating polarized wBL for 3 h at 22°C, chloroplasts localized along the anticlinal wall as the accumulation response (Fig 2C). When the cells were transferred to 5°C under the same polarized wBL condition, chloroplasts relocated from the anticlinal wall to the periclinal wall, indicating that chloroplasts avoided the polarized wBL at 5°C. These results confirm that the cold-avoidance response is due to avoidance from wBL under cold conditions, which is consistent with the microbeam analysis in our previous study [6].

**Fig 2.**
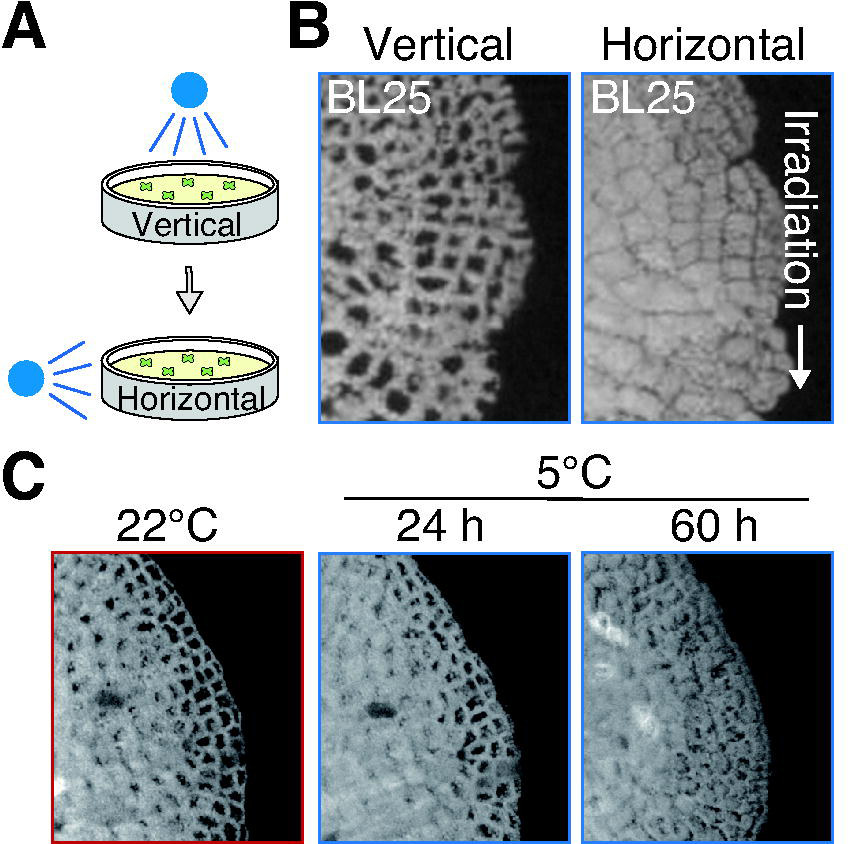
Chloroplasts avoid weak blue light under cold conditions. (A) Diagram of the experimental conditions used to induce the cold-avoidance response with vertical or horizontal irradiation of blue-light. After incubation under 25 µmol photons m^-2^ s^-1^ of blue-light (BL25) with vertical irradiation at 5°C for 24 h, the WT gemmalings were irradiated by horizontal BL25. (B) Chlorophyll fluorescence images after irradiation of vertical (left) or horizontal (right) BL as shown in (A). The white arrow indicates the light direction. (C) Chlorophyll fluorescence images after irradiation of the polarized weak BL. Ten-day-old WT sporelings were irradiated by vertically vibrating polarized weak BL at 22°C. The cells were transferred to 5°C, followed by incubation for 24 h and 60 h.

### Cold treatment decreases the transitional light intensity between the accumulation and avoidance responses

We hypothesized that the cold-avoidance response is induced by a cold-mediated decrease of the transitional light intensity (Fig 1). If this is the case, the accumulation response would be induced under much weaker BL in cold conditions. To investigate whether the accumulation response is induced at 5°C, WT cells were incubated under different intensities of wBL (5, 10, 15 and 25 µmol photons m^-2^ s^-1^; BL5, BL10, BL15 and BL25, respectively) (Fig 3A and 3B). Chloroplast position was quantified by a P/A ratio method, developed in earlier studies [4, 5]. The P/A ratio is an arbitrary unit calculated by dividing the chlorophyll fluorescence intensity along the periclinal cell wall by that along the anticlinal cell wall, with a higher value indicating periclinal positioning of chloroplasts and vice versa. In WT cells, the cold-avoidance response was observed at 5°C under BL10, BL15 and BL25 (Fig 3A), and at these higher BL intensity levels, we observed a decrease in P/A ratios (Fig 3B). However, the cold-avoidance response was not observed at 5°C under BL5 (Fig 3A) and the P/A ratio was unchanged during the incubation of these samples (Fig 3B). These results suggest that BL5 induces the accumulation response at 5°C. However, because similar chloroplast positioning was observed under dark conditions at 5°C [6], it is also possible that the cells were unable to respond to the extremely weak intensity of BL5. To clarify whether the accumulation response is induced under BL5 or not, an additional test was carried out. Following the induction of the cold-avoidance response at 5°C for 24 h under BL25, the light intensity was changed to BL5 or dark conditions (Fig 3C). After incubation for 24 h, BL5 conditions induced the accumulation response at 5°C, whereas under dark conditions there was no change in chloroplast position (Fig 3D). These results indicate that the accumulation response is induced under BL5 at 5°C. Taken together, these results indicate that the transitional light intensity is modulated by the temperature change (Fig 1).

**Fig 3.**
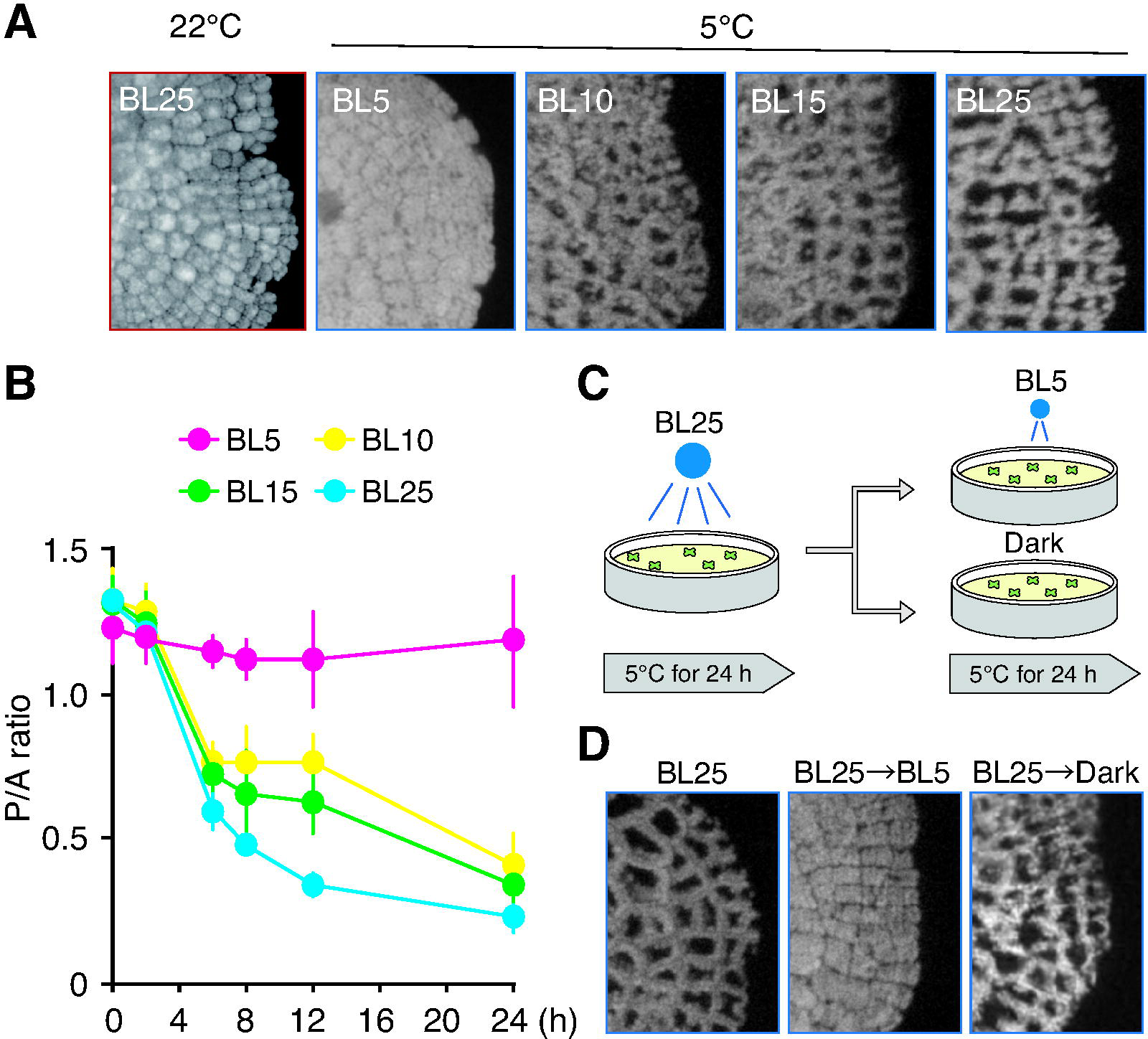
Cold-induced reduction of the transitional light intensity. (A) Chloroplast positioning in WT gemmalings incubated at 5°C for 12 h with 5, 10, 15, and 25 µmol photons m^-2^ s^-1^ of blue-light (BL5, BL10, BL15, and BL25, respectively). After pre-incubation of 1-day-old WT gemmalings under BL25 at 22°C for 1 h to induce the accumulation response, the cells were transferred to the indicated BL conditions at 5°C. (B) The P/A ratios, calculated from chlorophyll fluorescence images after incubation for 0, 2, 6, 8, 12, and 24 h. Error bars indicate standard deviations. (C) Diagram of experimental conditions used to confirm the cold-induced reduction of the transitional intensity. After pre-incubation under BL25 at 5°C for 24 h, WT gemmalings were transferred to BL5 or dark and incubated at 5°C for 24 h. (D) Representative chlorophyll fluorescence images of WT gemmalings after pre-incubation (BL25) and incubation under BL5 (from BL25 to BL5) or dark (from BL25 to Dark) as shown in (C).

### Artificial overexpression of Mpphot in *M. polymorpha* induces the avoidance response under wBL without cold treatment

We next considered the possibility that changes in endogenous phot expression may play a role in the change in transitional light intensity, as previous studies using *A. capillus-veneris* and *A. thaliana* suggest that a reduction in the transitional light intensity can be the result of artificial overexpression of phot [11, 12]. In *A. capillus-veneris* at 25°C, transient overexpression of the Ac*PHOT2-GFP* gene induced the avoidance response under wBL [11]. Similarly, the avoidance response was observed under wBL in transgenic *A. thaliana* overexpressing the At*PHOT2-GFP* gene at 23°C [12]. Consistent with these previous studies, we found that artificial overexpression of Mpphot fused to the yellow fluorescent protein Citrine (Mpphot-Citrine) in WT cells (35S::Mpphot-Citrine/WT) induced the avoidance response under BL25 at 22°C (without cold treatment), and degree of the avoidance response was dependent on Mpphot-Citrine expression levels (Fig 4A-4C). In the overexpression line (OX2) under BL5, the accumulation response at 22°C and cold-avoidance response at 5°C could be induced (Fig 4D and 4E), indicating that the transitional intensity is reduced by overexpression of Mpphot-Citrine. Therefore, it is conceivable that increase of the endogenous amount of Mpphot might play a role in the induction of the cold-avoidance response.

**Fig 4.**
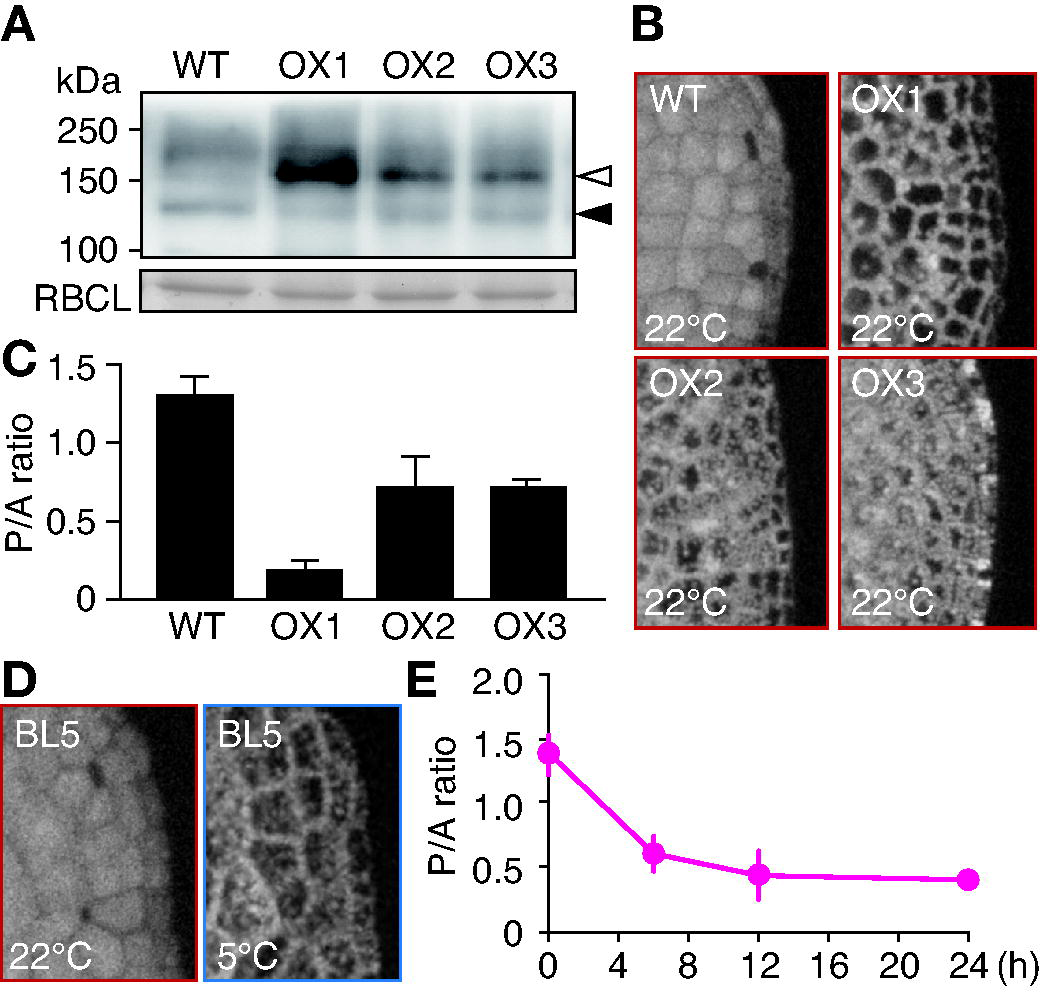
Overexpression of Mpphot-Citrine decreases the transitional light intensity. (A) Immunoblot analysis of WT and Mpphot-Citrine overexpression lines (35S::Mpphot-Citrine/WT: OX1, OX2, and OX3) with anti-Mpphot antibody. White and black arrowheads indicate Mpphot-Citrine and Mpphot, respectively. Rubisco large-subunit (RBCL) was shown as a loading control. (B) Representative images of chlorophyll fluorescence in WT and Mpphot-Citrine overexpression lines. (C) The P/A ratios calculated from the chlorophyll fluorescence images shown in (B) after the incubation under BL25 at 22°C for 12 h. (D) Confirmation of the reduction of transitional light intensity in OX2. In the 1-day-old gemmalings from OX2, the accumulation response was induced under BL5 at 22°C (left panel). The cold-avoidance response was induced under BL5 at 5°C for 24 h (right panel). (E) Time-course of the cold-avoidance response induced under BL5 at 5°C in OX2. The P/A ratios were calculated from the chlorophyll fluorescence images after incubation for 0, 6, 12, and 24 h.

### Alteration of endogenous Mpphot is not required for the induction of the cold-avoidance response

To examine the role of Mpphot expression in the induction of the cold-avoidance response, the endogenous amount of Mpphot was determined using immunoblot analysis. After gemmalings were transferred from the white light condition to BL25 condition at 22°C, the endogenous amount of Mpphot was increased at 6 h and the increased amount was maintained until 24 h (S Fig 1). To avoid the effect of the light condition change (i.e., from white light to BL25), the gemmalings were pre-incubated under BL25 at 22°C for 24 h. When temperature was changed from 22°C to 5°C, the endogenous amount of Mpphot was not changed at 3, 6, 9 h (Fig 5A). Note that this experimental condition was not able to separate phosphorylated and dephosphorylated forms of Mpphot. When we incubated the gemmalings under the same light and temperature condition as shown in Fig 5A, the cold-avoidance response was induced (Fig 5B), and the P/A ratio was decreased (Fig 5C). Unexpectedly the pre-incubation under BL25 at 22°C for 24 h increased the P/A ratio than the white light condition at 22°C (see 0 h in Fig 5C and 0 h in Fig 3B). Anyway, an increase of the endogenous amount of Mpphot is not required for induction of the cold-avoidance response in *M. polymorpha*. To confirm this result, we examined the complementation lines of Mpphot-Citrine under the control of the CaMV35S promoter with the genetic background of Mp*phot^KO^* (35S::Mpphot-Citrine/Mp*phot^KO^*), as CaMV35S promoter activity is reported to be suppressed by cold treatment in tobacco plants [24]. In transgenic *M. polymorpha* expressing Citrine under the control of the CaMV35S promoter (35S::Citrine/WT), we also confirmed that the promoter activity is slightly suppressed by 5°C treatment in *M. polymorpha* (Fig 5D). In the complementation lines (35S::Mpphot-Citrine/Mp*phot^KO^*), the cold-avoidance response was clearly observed (Fig 5E), confirming that the induction of the cold-avoidance response does not require any increase of Mpphot amount. Given that cold treatment increased the autophosphorylation status of phot [6, 25], we conclude that an increase of phot autophosphorylation status decreases the transitional light intensity without alteration of the endogenous amount of Mpphot.

**Fig 5.**
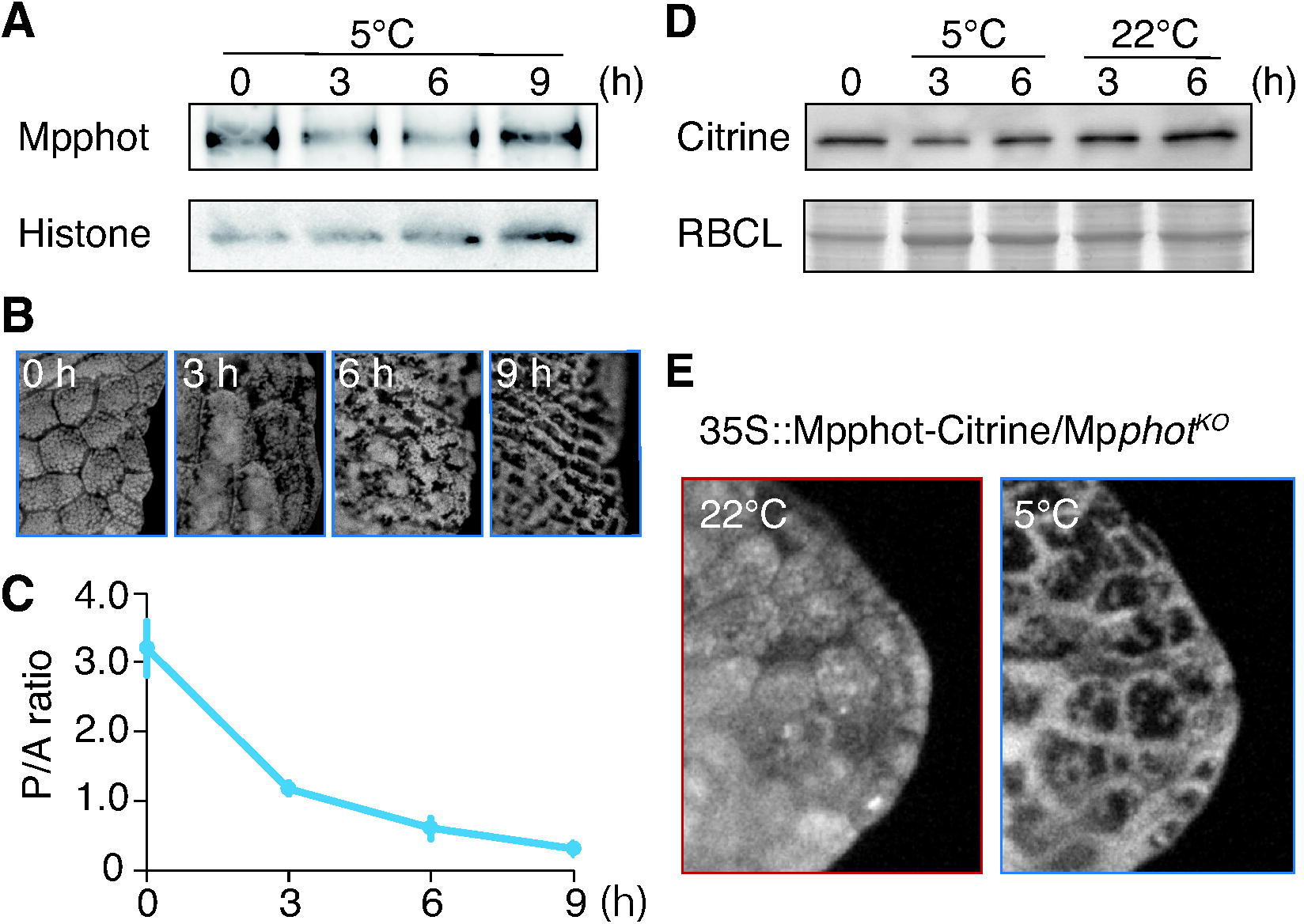
Alteration of Mpphot expression is not required for the induction of the cold-avoidance response. (A) Immunoblot analysis of endogenous MpPHOT amount in 4-day-old WT gemmaling (white light for 3 days and BL25 for 24 h at 22°C), incubated under BL25 at 5°C. The black arrowhead and asterisk indicate MpPHOT and non-specific signal, respectively. Histone H3 protein is shown as a loading control. (B) Representative chlorophyll fluorescence images of WT gemmalings incubated under the same light and temperature conditions as shown in (A). (C) The P/A ratios, calculated from chlorophyll fluorescence images after incubation for 0, 3, 6 and 9 h. Error bars indicate standard deviations. (D) Immunoblot analysis of cold-induced repression of activity of CaMV35S promoter. A transformant expressing Citrine under the control of CaMV35S promoter (35S::Citrine/WT) was used. Coomassie Brilliant Blue (CBB)-stained Rubisco large-subunit (RBCL) was used as a loading control. (E) Chlorophyll fluorescence images showing the cold-avoidance response in complementation lines (35S::Mpphot-Citrine/Mp*phot^KO^*). After incubation under BL25 at 22°C, the gemmalings were transferred to 5°C under the same light conditions.

## Discussion

### The transitional light intensity between the accumulation and avoidance responses can be changed

Light-induced chloroplast movement is widely found in photosynthetic organisms from algae to land plants [3]. The transitional light intensity between the accumulation and avoidance responses differs among various plant species [26]. Furthermore, the transitional light intensity can be reduced in response to low temperature, inducing the cold-avoidance response [4–6]. In this study, we analyzed the light fluence rate dependency of the cold-avoidance response. Under cold conditions, chloroplasts moved in a light fluence rate-dependent manner (Fig 3A and 3B), as occurs during the avoidance response at a standard growth temperature [27]. This result supports the conclusion that there is a cold-induced reduction of the transitional light intensity (Fig 1). Because the cold-avoidance response has been found to moderate photodamage in *M. polymorpha* [6], the plasticity of the transitional light intensity is vital for plants exposed to low temperatures. In *A. thaliana*, the transitional light intensity is controlled by two types of phot: Atphot1 and Atphot2. Atphot1 specifically regulates the accumulation response, while Atphot2 regulates both the accumulation and avoidance responses [7–9]. In this context, the coordination mechanism of the transitional light intensity in *A. thaliana* appears to be more complex than that in *M. polymorpha*. Further analysis of the transitional light intensity in various plant species and genetic mutants may lead to a better understanding of the transition mechanism of chloroplast relocation movements.

### Photoactivated phototropin amount changes the transitional light intensity

Intracellular phot proteins are more frequently activated under strong light than weak light conditions. Weak light irradiation at lower temperatures, prolongs the lifetime of photoactivated phot [6], and like the effect of strong light irradiation at a standard growth temperature, results in the increase of photoactivated phot population. Therefore, we concluded that the intracellular amount of photoactivated phot determines the transition between the accumulation and avoidance responses [6]. Furthermore, considering the effect of phot overexpression on the transitional light intensity (Fig 4), the transition mechanism may be based on the absolute amount of photoactivated phot in the cell rather than the ratio of active/inactive phot. How an increase in photoactivated phot proteins triggers the switch from the accumulation to avoidance responses remains unknown. Because phot autophosphorylation is also increased under strong-light/warm or weak-light/cold conditions [6,10,25], we speculate that to induce the avoidance response, an increase of photoactivated phot amount may change the phot phosphorylation status, possibly via intermolecular phosphorylation activity [28, 29]. Notably, the recruitment of phot to microdomains at the plasma membrane is involved in phot autophosphorylation [30]. Further autophosphorylation analysis regarding the spatial density of photoactivated phot may clarify the switching mechanism between the accumulation and avoidance responses.

### Alteration of phot expression is not required for induction of the cold-avoidance response in *M. polymorpha*

The cold-avoidance response is initiated by the temperature-dependent biochemistry of the phot LOV2 domain [6]. However, in *A. thaliana*, alteration of the endogenous amount of Atphot1 and Atphot2 in response to temperature was reported; lower temperature decreases Atphot1 expression and increases Atphot2 expression [16]. In this study, we showed that the cold-avoidance response can be induced regardless of the change in the endogenous Mpphot amount (Fig 5A-5C) and even without its promoter (Fig 5D and 5E), indicating that transcriptional regulation of Mpphot is not required for the cold-avoidance response in *M. polymorpha*. Mpphot is similar to Atphot2, which regulates both the accumulation and avoidance responses [17]. Consistent with Mpphot, Atphot2 expression in *A. thaliana* was reported to be unchanged at a temperature range from 4°C to 23°C [16]. Interestingly, cold treatment had a more significant impact on the decrease of Atphot1 expression than the increase of Atphot2 expression [16], implying that the expression of Atphot1 might be more sensitive to cold than that of Atphot2. The gene duplication of phot [31] and functional differentiation between phot1 and phot2 may have enabled land plants to have different transitional light intensities and chloroplast movements, allowing species to expand their range of habitat.

## Acknowledgements

The authors thank Dr. Takayuki Kohchi (Kyoto University) and Dr. Kimitsune Ishizaki (Kobe University) for providing *M. polymorpha* (WT and Mp*phot^KO^*) and the binary vectors (pMpGWB106 and pMpGWB306). The authors also thank Ms. Lee Kien Yong (Utsunomiya University) for helping the preparation of the manuscript. This work was supported by the Japan Society for the Promotion of Science KAKENHI [grant No. 18H02455 to Y.K.]; the Japan Science and Technology Agency, Exploratory Research for Advanced Technology, Numata Organelle Reaction Cluster [grant No. JPMJER1602 to Y.K.]; and the Plant Transgenic Design Initiative of the University of Tsukuba to Y.K.

## Figure Legends

**S Fig 1. Effect of the light condition change on the endogenous Mpphot expression.** Immunoblot analysis of endogenous Mpphot amount in WT gemmaling, incubated under BL25 at 22°C after culture under the white light condition (75 µmol photons m^-2^s^-1^) at 22°C for 3 days. The black arrowhead and asterisk indicate Mppot and non-specific signal, respectively. Histone H3 protein is shown as a loading control.

## References

1. Gotoh E, Suetsugu N, Yamori W, Ishishita K, Kiyabu R, Fukuda M, et al. Chloroplast accumulation response enhances leaf photosynthesis and plant biomass production. Plant Physiol. 2018;178: 1358–1369. doi:10.1104/pp.18.00484

2. Kasahara M, Kagawa T, Oikawa K, Suetsugu N, Miyao M, Wada M. Chloroplast avoidance movement reduces photodamage in plants. Nature. 2002;420: 829–832. doi:10.1038/nature01202.1.

3. Senn G. Die gestalts-und lageveränderung der pflanzen-chromatophoren. Leipzig: Wilhelm-Engelmann; 1908.

4. Kodama Y, Tsuboi H, Kagawa T, Wada M. Low temperature-induced chloroplast relocation mediated by a blue light receptor, phototropin 2, in fern gametophytes. J Plant Res. 2008;121: 441–448. doi:10.1007/s10265-008-0165-9

5. Ogasawara Y, Ishizaki K, Kohchi T, Kodama Y. Cold-induced organelle relocation in the liverwort Marchantia polymorpha L. Plant, Cell Environ. 2013;36: 1520–1528. doi:10.1111/pce.12085

6. Fujii Y, Tanaka H, Konno N, Ogasawara Y, Hamashima N, Tamura S, et al. Phototropin perceives temperature based on the lifetime of its photoactivated state. Proc Natl Acad Sci U S A. 2017;114: 9206–9211. doi:10.1073/pnas.1704462114

7. Jarillo JA, Gabrys H, Capel J, Alonso JM, Ecker JR, Cashmore a R. Phototropin-related NPL1 controls chloroplast relocation induced by blue light. Nature. 2001;410: 952–954. doi:10.1038/35073622

8. Kagawa T, Sakai T, Suetsugu N, Oikawa K, Ishiguro S, Kato T, et al. Arabidopsis NPL1: a phototropin homolog controlling the chloroplast high-light avoidance response. Science. 2001;291: 2138–2141. doi:10.1126/science.291.5511.2138

9. Sakai T, Kagawa T, Kasahara M, Swartz TE, Christie JM, Briggs WR, et al. Arabidopsis nph1 and npl1: blue light receptors that mediate both phototropism and chloroplast relocation. Proc Natl Acad Sci U S A. 2001;98: 6969–6974. doi:10.1073/pnas.101137598

10. Inoue S-I, Kinoshita T, Matsumoto M, Nakayama KI, Doi M, Shimazaki K-I. Blue light-induced autophosphorylation of phototropin is a primary step for signaling. Proc Natl Acad Sci U S A. 2008;105: 5626–5631. doi:10.1073/pnas.0709189105

11. Kagawa T, Kasahara M, Abe T, Yoshida S, Wada M. Function analysis of phototropin2 using fern mutants deficient in blue light-induced chloroplast avoidance movement. Plant Cell Physiol. 2004;45: 416–426. doi:10.1093/pcp/pch045

12. Kong SG, Kinoshita T, Shimazaki KI, Mochizuki N, Suzuki T, Nagatani A. The C-terminal kinase fragment of Arabidopsis phototropin 2 triggers constitutive phototropin responses. Plant J. 2007;51: 862–873. doi:10.1111/j.1365-313X.2007.03187.x

13. Kanegae H, Tahir M, Savazzini F, Yamamoto K, Yano M, Sasaki T, et al. Rice NPH1 homologues, OsNPH1a and OsNPH1b, are differently photoregulated. Plant Cell Physiol. 2000;41: 415–423. doi:10.1093/pcp/41.4.415

14. Jain M, Sharma P, Tyagi SB, Tyagi AK, Khurana JP. Light regulation and differential tissue-specific expression of phototropin homologues from rice (Oryza sativa ssp. indica). Plant Sci. 2007;172: 164–171. doi:10.1016/j.plantsci.2006.08.003

15. Suzuki H, Okamoto A, Kojima A, Nishimura T, Takano M, Kagawa T, et al. Blue-light regulation of ZmPHOT1 and ZmPHOT2 gene expression and the possible involvement of Zmphot1 in phototropism in maize coleoptiles. Planta. 2014;240: 251–261. doi:10.1007/s00425-014-2082-6

16. Łabuz J, Hermanowicz P, Gabry H. The impact of temperature on blue light induced chloroplast movements i^ś^n Arabidopsis thaliana. Plant Sci. 2015;239: 238–249. doi:10.1016/j.plantsci.2015.07.013

17. Komatsu A, Terai M, Ishizaki K, Suetsugu N, Tsuboi H, Nishihama R, et al. Phototropin encoded by a single-copy gene mediates chloroplast photorelocation movements in the liverwort Marchantia polymorpha. Plant Physiol. 2014;166: 411–427. doi:10.1104/pp.114.245100

18. Ishizaki K, Nishihama R, Ueda M, Inoue K, Ishida S, Nishimura Y, et al. Development of gateway binary vector series with four different selection markers for the liverwort Marchantia polymorpha. PLoS One. 2015;10: e0138876. doi:10.1371/journal.pone.0138876

19. Kodama Y. Time gating of chloroplast autofluorescence allows clearer fluorescence imaging in planta. Abraham T, editor. PLoS One. 2016;11: e0152484. doi:10.1371/journal.pone.0152484

20. Tsuboyama-Tanaka S, Kodama Y. AgarTrap-mediated genetic transformation using intact gemmae/gemmalings of the liverwort Marchantia polymorpha L. J Plant Res. 2015;128: 337–344. doi:10.1007/s10265-014-0695-2

21. Tsuboyama S, Kodama Y. AgarTrap protocols on your benchtop: Simple methods for Agrobacterium-mediated genetic transformation of the liverwort Marchantia polymorpha. Plant Biotechnol. 2018; in press.

22. Yatsuhashi H, Kadota A, Wada M. Blue- and red-light action in photoorientation of chloroplasts in Adiantum protonemata. Planta. 1985;165: 43–50. doi:10.1007/BF00392210

23. Kagawa T, Wada M. Polarized light induces nuclear migration in prothallial cells of Adiantum capillus-veneris L. Planta An Int J Plant Biol. 1995;196: 775–780. doi:10.1007/BF01106773

24. Schnurr JA, Guerra DJ. The CaMV-35S promoter is sensitive to shortened photoperiod in transgenic tobacco. Plant Cell Rep. 2000;19: 279–282. doi:10.1007/s002990050012

25. Okajima K, Kashojiya S, Tokutomi S. Photosensitivity of kinase activation by blue light involves the lifetime of a cysteinyl-flavin adduct intermediate, S390, in the photoreaction cycle of the LOV2 domain in phototropin, a plant blue light receptor. J Biol Chem. 2012;287: 40972–40981. doi:10.1074/jbc.M112.406512

26. Königer M, Bollinger N. Chloroplast movement behavior varies widely among species and does not correlate with high light stress tolerance. Planta. 2012;236: 411–426. doi:10.1007/s00425-012-1619-9

27. Kagawa T, Wada M. Chloroplast-avoidance response induced by high-fluence blue light in prothallial cells of the fern Adiantum capillus-veneris as analyzed by microbeam irradiation. Plant Physiol. 1999;119: 917–923. doi:10.1104/pp.119.3.917

28. Cho HY, Tseng TS, Kaiserli E, Sullivan S, Christie JM, Briggs WR. Physiological roles of the light, oxygen, or voltage domains of phototropin 1 and phototropin 2 in arabidopsis. Plant Physiol. 2007;143: 517–529. doi:10.1104/pp.106.089839

29. Kaiserli E, Sullivan S, Jones MA, Feeney KA, Christie JM. Domain swapping to assess the mechanistic basis of Arabidopsis phototropin 1 receptor kinase activation and endocytosis by blue light. Plant Cell. 2009;21: 3226–3244. doi:10.1105/tpc.109.067876

30. Xue Y, Xing J, Wan Y, Lv X, Fan L, Zhang Y, et al. Arabidopsis Blue Light Receptor Phototropin 1 Undergoes Blue Light-Induced Activation in Membrane Microdomains. Mol Plant. 2018;11: 846–859. doi:10.1016/j.molp.2018.04.003

31. Li FW, Rothfels CJ, Melkonian M, Villarreal JC, Stevenson DW, Graham SW, et al. The origin and evolution of phototropins. Front Plant Sci. 2015;6: 1–11. doi:10.3389/fpls.2015.00637

