## Supplemental Figure 1 for "Cold-induced switch in direction of chloroplast relocation occurs independently of changes in endogenous phototropin expression"

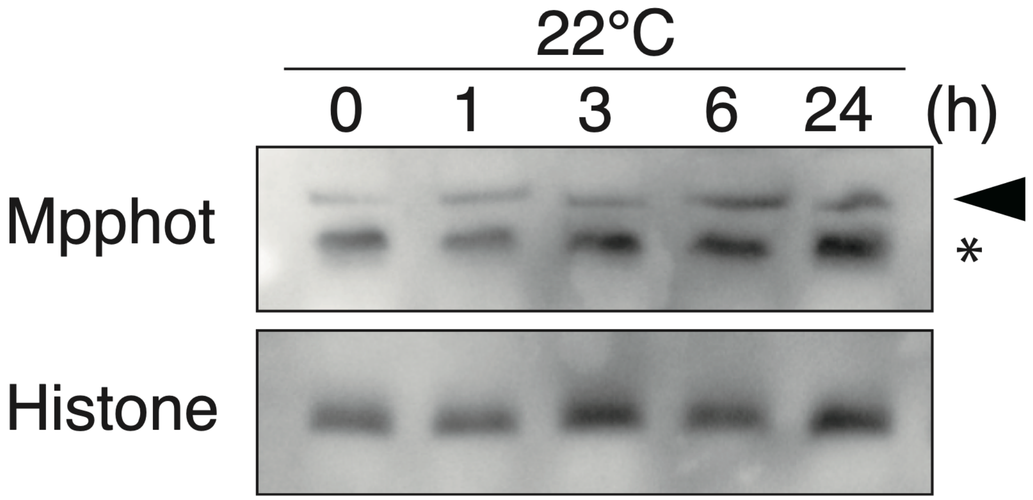


**S Fig 1. Effect of the light condition change on the endogenous Mpphot expression.** Immunoblot analysis of endogenous Mpphot amount in WT gemmaling, incubated under BL25 at 22° C after culture under the white light condition (75 µmol photons $m^{-2}$ $s^{-1}$) at 22° C for 3 days. The black arrowhead and asterisk indicate Mppot and non-specific signal, respectively. Histone H3 protein is shown as a loading control.
